# Ovalbumin-loaded mesoporous silica nanoparticles for allergen specific immunotherapy

**DOI:** 10.1101/2024.12.26.630362

**Authors:** Ana M. Pérez-Moreno, Pablo Torres, María del Carmen Martín-Astorga, Paula Cuevas-Delgado, Irene García-Esteban, José A. Céspedes, María I. Montañez, María José Torres, Carlos J. Aranda, Cristobalina Mayorga, Juan L. Paris

## Abstract

Allergic diseases are caused by an unnecessary immune response against harmless external substances (allergens), and they pose an important economic burden for healthcare systems with a large impact on the quality of life of patients. Allergen-specific immunotherapy (AIT) is the only treatment option capable of modifying the natural history of the disease, but current AIT schemes present safety and efficacy limitations. One possible strategy to address these limitations is to encapsulate the allergen in nanoparticle carriers that can deliver it to antigen presenting cells while hiding it from effector cells responsible for the allergic reaction. In this work, we evaluate the use of allergen-loaded mesoporous silica nanoparticles (MSNs) as AIT agents. MSNs of different pore sizes were prepared and characterized, evaluating their capacity to load and release ovalbumin (OVA) as a model allergen. Extra-large pore MSNs (XL-MSNs) showed the optimal loading and release behavior, presenting also enhanced activation of the dendritic cell line DC2.4 and reduced allergenic capacity in pre-sensitized RBL- 2H3 cells, both compared to free OVA. After evaluating their biodistribution following subcutaneous, sublingual or intravenous administration, their therapeutic potential in AIT was further assessed in an *in vivo* murine model of OVA systemic anaphylaxis. The results showed that intravenous administration of OVA-loaded XL-MSNs significantly protected the mice from anaphylaxis and induced a Th1/Treg-immune profile, while administration through other routes failed to prevent the development of an anaphylactic reaction upon provocation with OVA. These findings establish MSNs, particularly via intravenous administration, as a promising platform to develop safer and more effective AIT.

## 1. Introduction

Allergic diseases are caused by an unnecessary immune response against harmless external substances (allergens) [1]. The incidence of allergies is increasing worldwide, becoming a growing economic burden for healthcare systems and having a large impact on the quality of life of patients [2]. The most common type of allergic reactions are immediate hypersensitivity reactions [1], which are associated with a T helper 2 (Th2) response. Specific immunoglobulin E (sIgE) against the allergen [3] binds to high-affinity receptors (FcεRI) on mast cells and basophils. When a sensitized individual is re-exposed to the allergen, the crosslinking of sIgE bound to the effector cell membrane triggers its activation, releasing histamine and other mediators that are responsible for the symptoms of the allergic reaction [4]. Currently, the management of allergic diseases is based on the avoidance of triggering allergens or symptom management using drugs such as antihistamines, corticosteroids, and adrenaline [5]. However, accidental exposure to the allergen will continue to trigger allergic reactions, which greatly affects the quality of life of allergic patients. Allergen-specific immunotherapy (AIT) is currently the only treatment option that can modify the natural history of the disease and induce long-term allergen tolerance by switching the immune response from a Th2 profile to a Th1 or regulatory (Treg) response [6]. In AIT, the allergen is repeatedly administered at a controlled dose to generate specific immune tolerance. Despite the good potential for AIT, safety and efficacy limitations may hinder its clinical utility.

As safety concerns arise from the possibility of triggering an allergic reaction using the responsible allergenic protein during AIT, one interesting approach is to encapsulate the allergen in a carrier formulation, usually a micro- or nanoparticle that provides a sustained release of the allergen [5,7–10]. This encapsulation limits the amount of free allergen available immediately after administration [11], reducing the likelihood of an undesired allergic reaction [12], while ensuring that enough allergen will be released at longer time points to provide therapeutic efficacy. Importantly, nanoparticles present additional advantages in AIT, such as their effect on the biodistribution of the therapeutic agent and their size-dependent interactions with immune cells. Regarding biodistribution, the nanoparticle size and surface characteristics as well as the route of administration, will determine the dose of nanoparticles that reach different tissues and organs, which in turn determines their therapeutic efficacy and safety profile [13]. In this regard, nanomaterial-based AIT approaches have been described for subcutaneous immunotherapy (SCIT) [14], sublingual immunotherapy (SLIT) [6] and intravenous immunotherapy (IVIT) [15–17]. SCIT is generally considered a more effective route of administration for AIT, although it presents a larger risk of triggering an allergic reaction than SLIT, which has a better safety profile but requires larger doses to produce benefitial effect [13]. It is important to mention that, while SCIT and SLIT are conventional routes of administration for AIT, IVIT is directly enabled by the biodistribution of systemically administered nanoparticles, which tend to accumulate in the spleen and liver, organs with large populations of antigen presenting cells (APCs), essentials for inducing an immune response. Once the nanoparticles have reached APCs such as dendritic cells [18–20], either in the subcutaneous region, in the sublingual mucosa, or in the liver or spleen, they are easily taken up by these cells, releasing a significant fraction of the therapeutic cargo intracellularly and potentially enhancing efficacy.

Most of the previous work on nanocarriers for AIT is based on the use of polymeric nanoparticles (such as PLGA [16,21–24], dendrimers [11] or Gantrez[25,26]), and some very promising results have been reported not only in *in vivo* animal models but even reaching clinical trials for some of these formulations[27]. However, there is a wide range of nanoparticle compositions that have remained largely unexplored for this application. In particular, mesoporous silica nanoparticles (MSNs) have been thoroughly evaluated for drug delivery in different diseases [28,29], but not for their application in AIT [30]. MSNs are biocompatible, and they are known to undergo dissolution under physiological conditions, giving rise to nontoxic products that can be safely excreted [31,32]. MSNs present a large surface area that provides high cargo loading capacity [28], and their pore size can be finely tuned to be adapted for specific cargo molecules or therapeutic applications [33–35]. Furthermore, the pore size of protein-loaded MSNs has been previously reported to determine their antigen delivery performance in cancer immunotherapy [36,37], and MSNs have also been suggested to have an inherent adjuvant effect [38]. Thus, in this work we aim to evaluate the potential of allergen-loaded MSNs of optimized pore sizes as therapeutic agents in allergic diseases. We prepared MSNs of different pore sizes, evaluating their cargo loading and release profile using ovalbumin (OVA) as a model allergen. We evaluated the immunogenicity and allergenicity of OVA-loaded MSNs through several *in vitro* and *in vivo* methods, and finally we studied the biodistribution and therapeutic efficacy *in vivo* in a mouse model of systemic anaphylaxis after administration through different routes (SCIT, SLIT, and IVIT).

## 2. Experimental section

### 2.1 Materials

The following reagents were purchased from Merck (Sigma‒Aldrich, Spain) and were used without further purification: tetraethylorthosilicate (TEOS), cyclohexane, triethanolamine, cetyltrimethylammonium chloride (CTAC), ammonium nitrate, ethanol, hydrochloric acid, fluorescein isothiocyanate (FITC), aminopropyltriethoxysilane (APTES), OVA, phosphate buffered saline (PBS) tablets, Roswell Park Memorial Institute (RPMI)-1640 culture medium, Eagle’s Minimum Essential Medium, foetal bovine serum (FBS), nonessential amino acids, L-glutamine, β-mercaptoethanol, cell viability/proliferation kit WST-1. Mouse anti-OVA IgE (Clone 2C6) was purchased from Bio-Rad, Spain.

AlexaFluor^TM^750 carboxylic acid, succinimidyl ester (AlexaFluor750-NHS), antibodies and other reagents for ELISA experiments were obtained from Thermo Fisher Scientific (Spain). Antibodies for flow cytometry were purchased from Biolegend, BD Biosciences or Invitrogen (Spain) according to Table S1.

### 2.2 Synthesis of MSNs

MSNs of different pore sizes were prepared by a previously described biphasic method based on the condensation of TEOS in a biphasic water/cyclohexane system, using triethanolamine as the base and CTAC as the structure-directing agent surfactant [39,40]. The aqueous phase was composed of a mixture of 24 mL of a commercial aqueous solution of CTAC (25% w/v)), 0.18 g of triethanolamine and 36 mL of deionized water. The organic phase consisted of 20 mL of a mixture of cyclohexane with TEOS. The concentration of TEOS depended on the material to be prepared: 40% for small-pore MSN (S-MSN), 20% for medium-pore MSN (M-MSN), 10% for large-pore MSN (L-MSN) and 5% for extra-large pore MSN (XL-MSN). The synthesis reaction was carried out at 50°C for 24 h. Then, the surfactant was extracted by ion exchange with an ethanolic solution of ammonium nitrate (10 mg/mL) at reflux for 1 h, followed by a second reflux for 2 h in an ethanolic solution of 12 mM HCl. Finally, the material was washed with ethanol 3 times to obtain the desired materials, which were dried and stored at room temperature until further use. Fluorescent MSN were also obtained by adding a pre-reacted mixture of 1.5 mg of fluorescein FITC or 500 µg of AlexaFluor750-NHS and 1.5 µL of APTES in 1 mL of ethanol in the aqueous phase during MSN synthesis.

### 2.3 Cargo loading and release from MSNs

First, OVA was labelled with FITC for the cargo loading and release experiments. To do this, 0.5 mg of FITC dissolved in 500 µL of DMSO were added to 10 mL of a 2 mg/mL solution of OVA in PBS and the mixture was stirred at room temperature for 24 h protected from light. Then, OVA-FITC was purified using a 30 kDa Amicon filter, washing the product first with PBS and finally with water until all nonreacted FITC was removed, obtaining OVA-FITC by freeze-drying. Non-labelled or FITC-labelled OVA (as a model allergen) was loaded in MSNs by dispersing 10 mg of MSNs in a 5 mg/mL protein solution in PBS (10 mM, pH 7.4) and stirring overnight. Then, the loaded particles were collected by centrifugation, and the non-loaded cargo was quantified from the supernatant by UV‒Vis spectrophotometry. For release experiments, 10 mg of OVA-FITC-loaded MSNs of different pore sizes were dispersed in 1.8 mL of PBS and 0.5 mL of MSN suspension were placed on a Transwell permeable support with 0.4 μm of polycarbonate membrane (3 replicas were performed). The well was filled with 1.5 mL of PBS and the suspension was stirred at 37 °C during all the experiment. At every time point studied, the solution outside the transwell insert was replaced with fresh PBS and the amount of cargo released was determined by fluorescence spectrometry (λ_EX_=480 nm; λ_EM_=515 nm).

### 2.4 Characterization techniques

Dynamic light scattering (DLS) and Z-potential measurements were performed with a Malvern Zetasizer Nano ZS90 instrument, checking both particle size and surface charge. The instrument used was equipped with a “red laser” (ʎ = 300 nm), and DLS measurements were performed with a detection angle of 90°, while the Smoluchowski approximation was used for Z-potential measurements. To check the morphology and the different pore sizes of the nanoparticles, the characterization of the nanoparticles was performed by transmission electron microscopy (TEM) on a Thermo Fisher Scientific Tecnai G2 20 Twin using copper grids of mesh size 200 coated with a Formvar-Carbon film. Nitrogen adsorption (in a Micromeritics ASAP 2020 unit) measurements were carried out at the Central Research Support Services (SCAI) of the University of Malaga (UMA). The pore width of the different MSN formulations was determined from the maximum Barrett-Joyner-Halenda (BJH) Adsorption Cumulative Pore Volume distribution *vs* pore width graph obtained by N_2_ adsorption. Fluorescence microscopy was carried out on a Nikon Eclipse Ti Fluorescence Microscope (Nikon, Japan). *In vivo* fluorescence was evaluated with In-Vivo Xtreme equipment (Bruker, Germany).

### 2.5 *In vitro* evaluation of MSNs in DC2.4 cells

A mouse dendritic cell line (DC 2.4) was used to evaluate the immunological effect of MSNs [19,37]. The day prior to the experiment, 50,000 DC2.4 cells were seeded in each well, using a 96 well plate. For cellular uptake and viability/proliferation experiments, DC 2.4 cells were incubated with FITC-labelled MSNs for 2 h at a concentration of 10 µg/mL (in complete culture medium, RPMI- 1640 supplemented with 10% fetal bovine serum, nonessential amino acids, L-glutamine and β-mercaptoethanol, as recommended by the distributor (Sigma‒ Aldrich)) at 37°C and 5% CO_2_. Twenty-four hours later, nanoparticle uptake was evaluated by flow cytometry and fluorescence microscopy. Cell viability/proliferation was evaluated using the commercial kit WST-1 following the manufacturer’s instructions. To evaluate the immunogenic effect, changes in the expression of CD40 (a marker of dendritic cell activation) were assessed by flow cytometry after incubation with empty and OVA-loaded nanoparticles (non-labelled).

### 2.6 *In vitro* evaluation of allergenic and viability capacity of MSNs in RBL-2H3 cells

*In vitro* allergenicity of OVA-loaded MSNs was assessed using the pre-sensitized basophil-like cell line RBL-2H3 by evaluating their degranulation through the measurement of β-hexosaminidase released. Briefly, RBL-2H3 cells were seeded in a 96 well plate in complete culture medium (Eagle’s Minimum Essential Medium supplemented with 15% fetal bovine serum, nonessential amino acids and L-glutamine) at 37°C and 5% CO_2_ and three hours later they were sensitized with mouse anti-OVA IgE at a final concentration of 0.1 µg/mL. Twenty-four hours after seeding, the cells were washed with Tyrode’s buffer (135 mM NaCl, 5 mM KCl, 1.8 mM CaCl_2_, 1.0 mM MgCl_2_, 5.6 mM Glucose, 20 mM HEPES, and 1 mg/ml BSA at pH 7.4). Then, the cells were incubated with empty MSNs, OVA-loaded MSNs or free OVA using equivalent concentrations of OVA, 10 µg/mL, in Tyrode’s buffer for 1 h at 37 °C. Then, the supernatants were collected, and the cells were lysed with 1% Triton X-100. Then, 50ul of each supernatant or lysate was mixed with 200ul of 1 mM p-nitrophenyl N-acetyl-beta-D-glucosamine in 0.05M citrate buffer (pH 4.5). After 1 h at 37 °C, the reaction was quenched with 500 µL of 0.05M sodium carbonate buffer (pH 10). The amount of β-hexosaminidase was determined by absorbance at 405 nm. β-hexosaminidase release for each condition was expressed as percentage of total enzyme in each sample (released to supernatant + cell lysates).

For cell viability determination under the different treatments, the same general procedure was followed up to the addition of p-nitrophenyl N-acetyl-beta-D-glucosamine, adding instead the substrate from the WST-1 cell viability/proliferation kit and following then the kit’s manufacturer’s instructions.

### 2.7 *In vivo* experiments in mice for biodistribution and immunological response assessement

Mouse studies were carried out following Spanish national and European regulations (RD1201/2005, 32/2007, 2010/63/UE and RD53/2013). The mice were hosted at IBIMA-Plataforma BIONAND (Registration No. ES 290670001687). All procedures followed the 3R principles and received appropriate regulatory approval before starting (protocol 18/11/2021/180 approved by both the Institutional Ethics Committee and by *Consejería de Agricultura, Ganadería, Pesca y Desarrollo Sostenible, Junta de Andalucía*). The mice were anaesthetized during the different procedures and finally sacrificed by cervical dislocation. After obtaining the corresponding samples, the mice were stored and incinerated according to institutional guidelines.

To evaluate the immunological effect of the nanoparticles *in vivo*, OVA-loaded XL- MSNs were administered subcutaneously once a week for 3 weeks. Five- to six-week-old BALB/c mice (male, n=3 mice per group) from Janvier Lab (Saint-Berthevin Cedex, France) were used. One week after the last administration, the mice were anaesthetized intraperitoneally, and blood was obtained from the retroorbital plexus before euthanizing the animals.

To evaluate the biodistribution of the nanoparticles after administration through different routes, Alexa750-labelled MSNs, fluorescent in the near infrared (NIR), were used. Using male mice (n=3), 150 µg of fluorescent MSN dispersed in PBS were administered either intravenously (IV, by tail vein or retro-orbital injection), subcutaneously (SC) or sublingually (SL). For IV and SC injection, the volume injected was 100 µL, while for SC administration, only 10 µL of MSN suspension were deposited with a pipette beneath the tongue of isoflurane-anesthetized animals. At different time points, nanoparticle fluorescence in the mice was examined using an *in vivo* imaging system, In-Vivo Xtreme (Bruker). At the endpoint (2 days after MSN administration), the mice were euthanized by cervical dislocation under anaesthesia and the different organs were also observed under the *in vivo* fluorescence imaging system.

To evaluate the therapeutic efficacy of OVA-loaded XL-MSNs as AIT agents, an OVA systemic anaphylaxis mouse model was generated. 5-6 weeks old Balb/c mice (n=8 mice per group, 4 male and 4 female) were used for these experiments. First, the mice were sensitized by intraperitoneally injecting them with 50 µg of OVA adsorbed in 2 mg of Alum once per week for 2 weeks. Then, mice were treated with OVA-loaded XL-MSNs administered as described for the biodistribution experiment weekly for 6 weeks (using retro-orbital injection for the IV route). Two control groups were also included, non-sensitized mice (healthy control) and sensitized mice without treatment (anaphylactic control). After the AIT scheme was complete, the therapeutic effect was evaluated by challenge of mice intraperitoneally with 500 µg of OVA. Symptoms were monitored (with a scoring system[16] from 0, normal behavior, to 5, animal death) and body temperature was measured rectally with a thermometer 20 min after challenge. Then, the animals were anesthetized intraperitoneally, and blood was collected retro-orbitally to obtain sera to measure specific antibodies by ELISA. Then the mice were euthanized by cervical dislocation and the lymph nodes (axillary, mandibular and mesenteric) and spleen were collected in RPMI containing 10 % FBS and mashed on a 40-μm cell strainer to prepare a single cell suspension.

### 2.8 Anti-OVA antibody detection by ELISA

Different specific anti-OVA antibodies (IgG1, IgG2a, IgG2b and IgE) were determined in the mouse sera by ELISA using biotinylated rat anti-mouse antibodies. For the ELISAs, high-binding ELISA 96-well plates were coated with OVA. After blocking the plate with a casein-containing buffer solution, the mouse sera were added (1:8 dilution for IgE detection, 1:50 dilution for IgG detection) and incubated overnight at 4°C. Then, biotinylated secondary antibodies were added, followed by the addition of avidin-horseradish peroxidase (HRP). Finally, the results were obtained by measuring the colorimetric conversion of an HRP substrate (TMB) after stopping the reaction with H_2_SO_4_ using a plate reader (λ_ABS_=450 nm). Thorough washing of the plate with PBS containing 0.05% Tween 20 was carried out between the different steps in the protocol.

### 2.9 Cellular immunological response evaluation

Red blood cells were lysed using 1x RBC Lysis Buffer (Invitrogen). Some of the obtained cells (2 x 10^6^ cells per well per condition in 48 well plates) were further cultured in complete culture medium with 2 µg/mL of OVA for 48 h and the supernatants were collected for released cytokine detection by ELISA. The rest of the cells were used for flow cytometry evaluation of the immune response. Before the surface staining, cells were incubated with LIVE/DEAD™ Fixable Blue (Invitrogen) for 15 minutes at room temperature and subsequently with TruStain FcX™ PLUS (anti-mouse CD16/32) antibody (Biolegend). After washing with a staining buffer (DPBS containing 2% FBS, 4mM EDTA and 0.02% of sodium azide), up to 5 x 10^6^ cells were suspended in a 100 μL staining antibody buffer at 4°C in the dark for 40 minutes containing surface antibodies as in **Table S1**. Cells were then washed twice with a staining buffer, fixed and permeabilized using Foxp3/Transcription Factor Staining Buffer Set (eBioscience) according to the manufacturer’s instructions. Intracellular staining was performed at 4°C overnight. Cell suspensions were washed 3 times in the Perm buffer before acquiring a Cytek™ 4-laser Aurora CS (Cytek Biosciences) cytometer. Results were analyzed using FlowJo software (Tree Star Inc., Ashland, OR). Gating strategy was performed as in **Figure S1**.

## 3. Results and Discussion

The successful preparation of MSNs of different pore sizes was confirmed by DLS, Z potential, TEM and N_2_ adsorption (**Figure 1**, **Table S2**).

**Figure 1.**
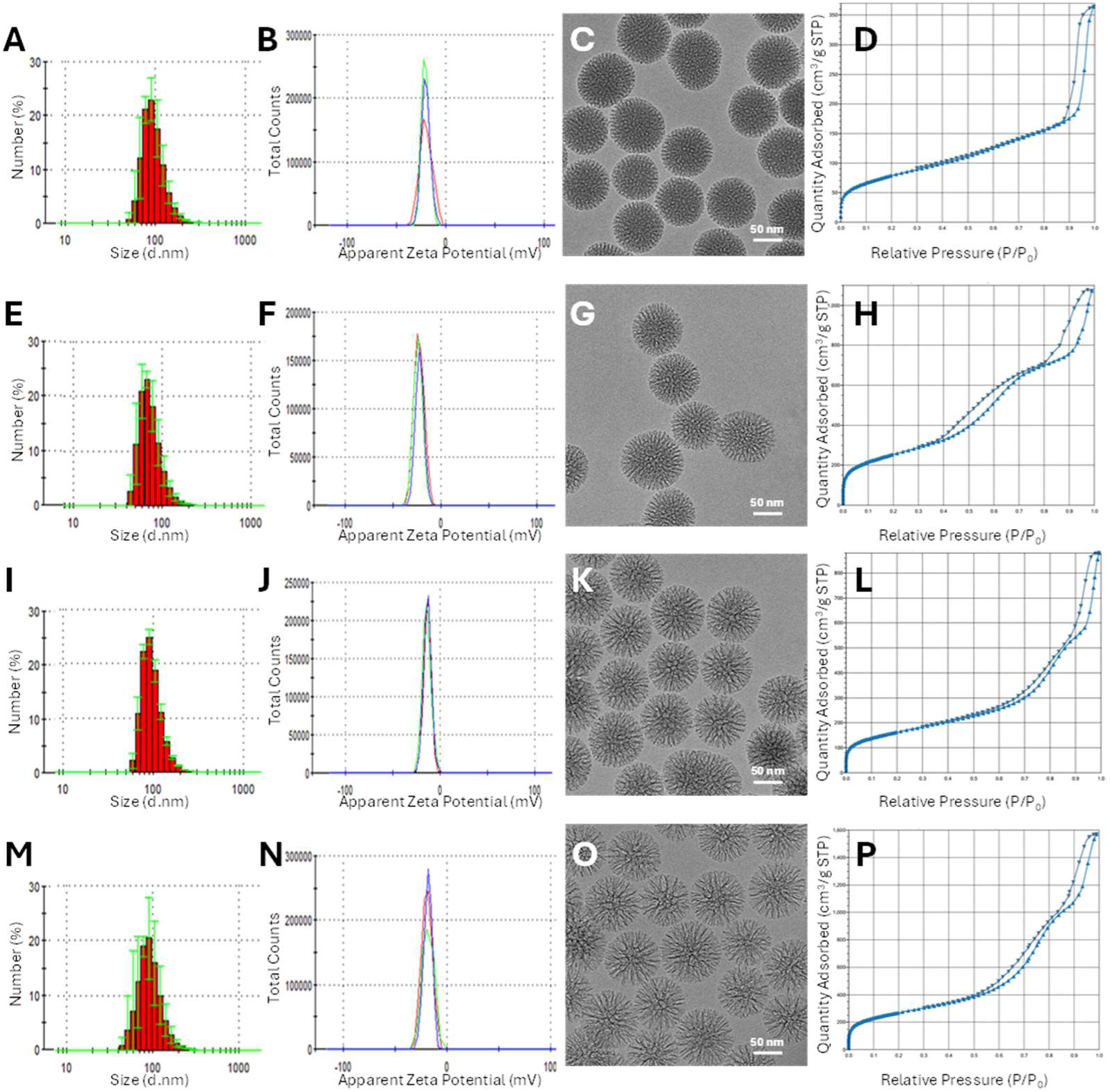
Characterization of S-MSN (A-D), M-MSN (E-H), L-MSN (I-L) and XL- MSN (M-P) by DLS (A,E,I,M), Zeta potential (B,F,J,N), TEM (C,G,K,O) and N_2_ adsorption (D,H,L,P).

DLS showed an adequate size distribution, with a Z average diameter of around 160 nm for all MSN types, which also presented negative Z potential, as a result of the presence of silanol groups on the nanoparticle surface after the synthesis and surfactant extraction (Table S2). All MSNs presented the expected porosity (**Figure 1**), with a large surface area (over 280 m^2^/g for all particle types) and tunable pore sizes: from 4.85 nm for S-MSN to 8.39 nm for XL-MSN (Table S2). These results are in line with previous reports on pore size-tunable MSNs prepared by the same biphasic method [34,35,40,41].

When evaluating the effect of MSN pore size on OVA loading, a direct relationship was found between pore diameter and protein loading (**Figure 2**), with XL-MSN loading larger amounts of OVA than the other evaluated materials, as their pores are likely too small to accommodate the protein inside. When analyzing OVA release, XL-MSN released a much larger amount than MSNs with smaller pores, probably due not only to the increased protein loading in this material, but also to the greater accessibility of the solvent to larger pores, facilitating cargo release. These results are in good agreement with previous reports, which had showed that MSNs with larger pores could load and release larger amounts of OVA [35,37,42], highlighting the potential of this material for antigen delivery.

**Figure 2.**
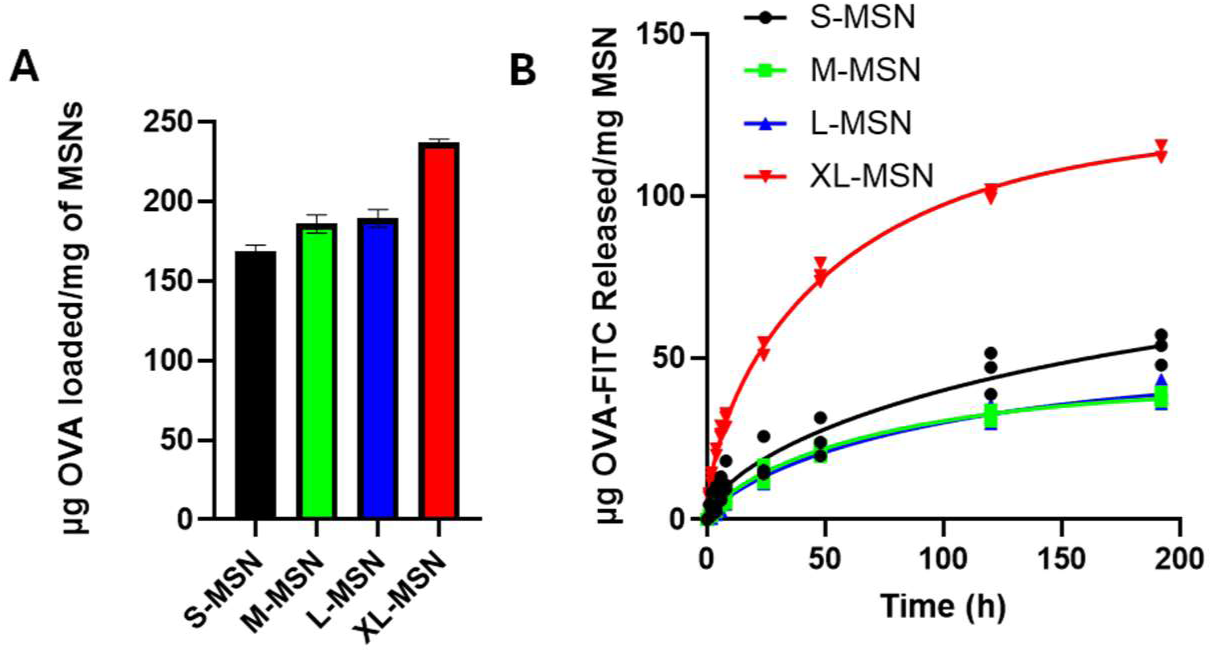
OVA-FITC loading (A) and release in PBS at 37 °C under continuous stirring (B) in/from MSNs of different pore sizes.

To evaluate the immunogenicity of OVA-loaded MSNs, an *in vitro* experiment with DC2.4 cells (a model dendritic cell line) was carried out first. The results (**Figure 3A-B**) show good biocompatibility of all MSN formulations up to 10 µg/mL, with some cytotoxicity observed at 20 µg/mL, especially for S-MSN. DC2.4 cells efficiently uptake all MSN types, although XL-MSNs presented a lower uptake percentage compared to the other types of MSNs (**Figure 3C**). When the activation of DC2.4 cells was assessed (by quantifying the percentage of cells expressing CD40 by flow cytometry), the results showed that OVA-loaded XL- MSNs induced the largest degree of dendritic cell activation, significantly decreasing the percentage of CD40^+^ cells as the nanoparticle pore size decreased (**Figure 3D**). Furthermore, treatment with empty MSNs or even with free OVA in the absence of a nanocarrier, even at high concentrations (up to 4 µg/mL), did not induce significant cell activation. These results are in good agreement with previous reports that showed that MSN pore size regulates their antigen delivery efficiency [37], with XL-MSNs loaded with OVA (similar to the those prepared in this work) being the optimal antigen delivery system and clearly enhancing the immunogenicity of the free protein.

**Figure 3.**
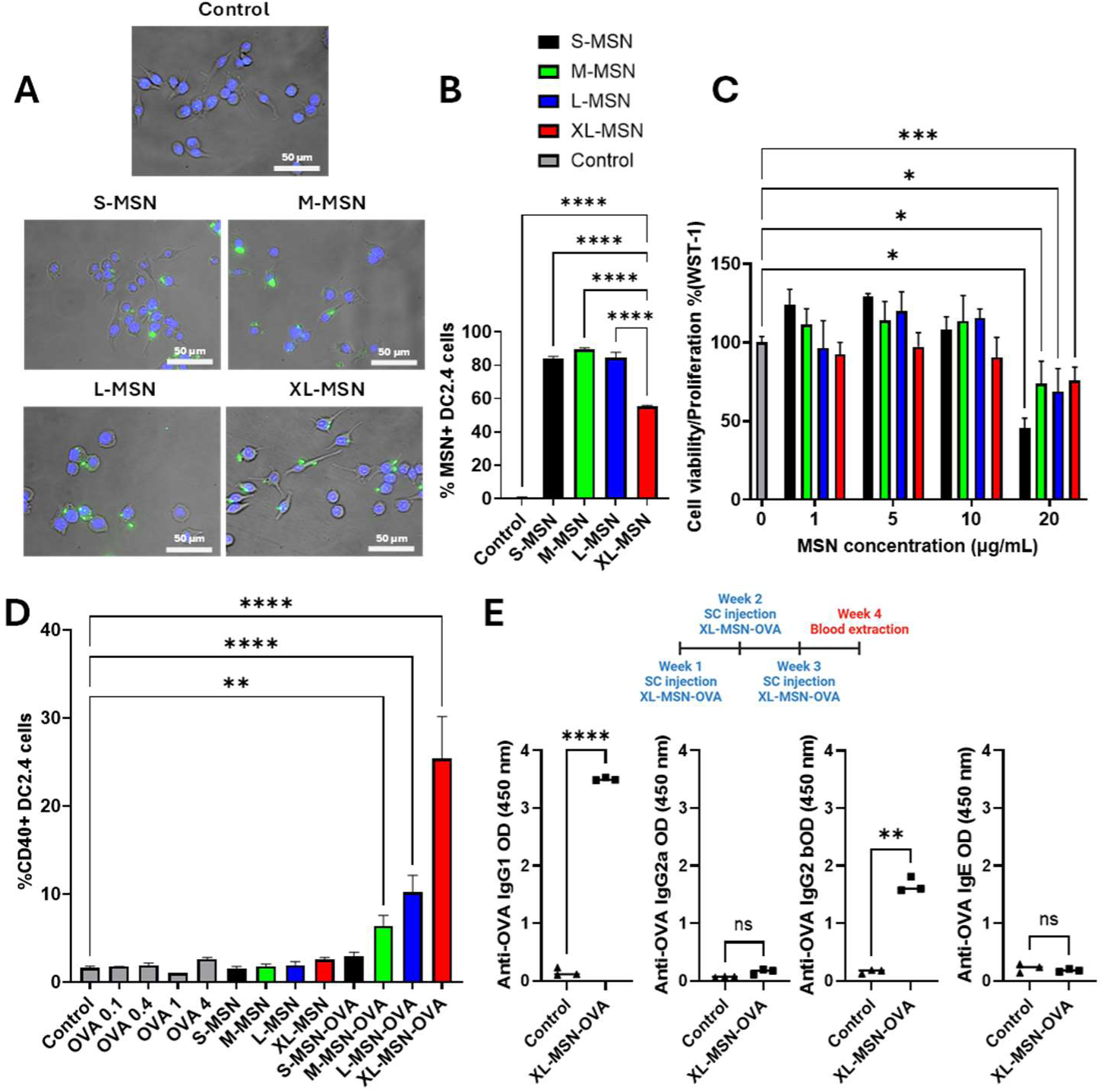
Immunogenicity evaluation of MSNs. *In vitro* evaluation of MSNs incubated with DC2.4 cells (A-D). Fluorescence microscopy images of DC2.4 cells incubated with different types of FITC-labeled MSNs at a concentration of 10 µg/mL. Blue fluorescence from cell nuclei (DAPI) and green fluorescence from MSNs (A); WST-1 assay results evaluating DC2.4 cell proliferation/viability after incubation with MSNs at different concentrations (B); Flow cytometry results showing the percentage of DC2.4 cells with nanoparticle uptake using different types of FITC-labeled MSNs at a concentration of 10 µg/mL (C); Activation of DC2.4 was evaluated by the expression of CD40 using flow cytometry upon treatment with free OVA (0.1-4 µg/mL), empty MSNs (10 µg/mL) and OVA-loaded MSNs (10 µg/mL) (D). *In vivo* immunization experiments in mice employing OVA-loaded XL-MSN subcutaneously injected 3 times weekly (E). OVA-specific antibodies (IgG1, IgG2a, IgG2b and IgE) were measured in sera by ELISA. Data are Means ±SD, n=3. Statistical analysis was performed using one way ANOVA. *p<0.05; **p<0.01; ***p<0.005; ****p<0.001.

To further evaluate the prospects of OVA-loaded XL-MSNs for immunization, these particles were subcutaneously administered three times (weekly) in BALB/c mice **(Figure 3E**). OVA-loaded XL-MSNs were used due to their higher capacity to activate DC2.4 cells. One week after the last administration, blood was obtained, and OVA-specific antibodies were evaluated in sera by ELISA. The repeated administration of OVA-loaded XL-MSN was well tolerated by the animals, with no perceivable changes in their behaviour and no significant differences in general serum biochemistry parameters (**Table S3**). The results of specific antibody generation (**Figure 3E**) showed a strong humoral response after three subcutaneous administrations of OVA-loaded XL-MSNs, with significant production of anti-OVA IgG1 and IgG2b compared to nontreated control mice. On the other hand, no significant differences were found between control and nanoparticle-treated mice in the production of specific anti-OVA IgG2a or IgE. These results highlight the promising nature of the developed platform for vaccination or immunotherapy.

Besides the need for improving AIT efficacy, the main factor limiting its clinical potential is the risk of triggering an allergic reaction during treatment. By using nanoparticles as allergen carriers, most of the allergen dose would initially be hidden from effector cells, potentially improving therapeutic safety. To evaluate this potential for MSNs, an *in vitro* experiment with pre-sensitized RBL-2H3 cells was first carried out. The results showed that MSNs were not toxic to RBL-2H3 cells (**Figure 4A**). When evaluating β-hexosaminidase release from pre-sensitized RBL-2H3 cells (**Figure 4B**), a significantly lower allergenic capacity of OVA-loaded MSNs compared to the same dose of free OVA was observed. Moreover, the largest decrease in ß-hexosaminidase release was observed with OVA-loaded XL-MSNs, which presented similar levels to the negative control (spontaneous release. The slightly larger β-hexosaminidase release percentage in OVA-loaded MSNs with smaller pore sizes could be due to the presence of OVA adsorbed on the nanoparticle surface, as the protein likely would not fit inside smaller pores, thereby being more exposed and available to interact with anti-OVA IgE on the cell membrane. As expected, empty MSNs also did not induce any β-hexosaminidase release. These results highlight the promising potential of allergen encapsulation inside MSNs, especially in XL-MSNs, to improve AIT safety.

**Figure 4.**
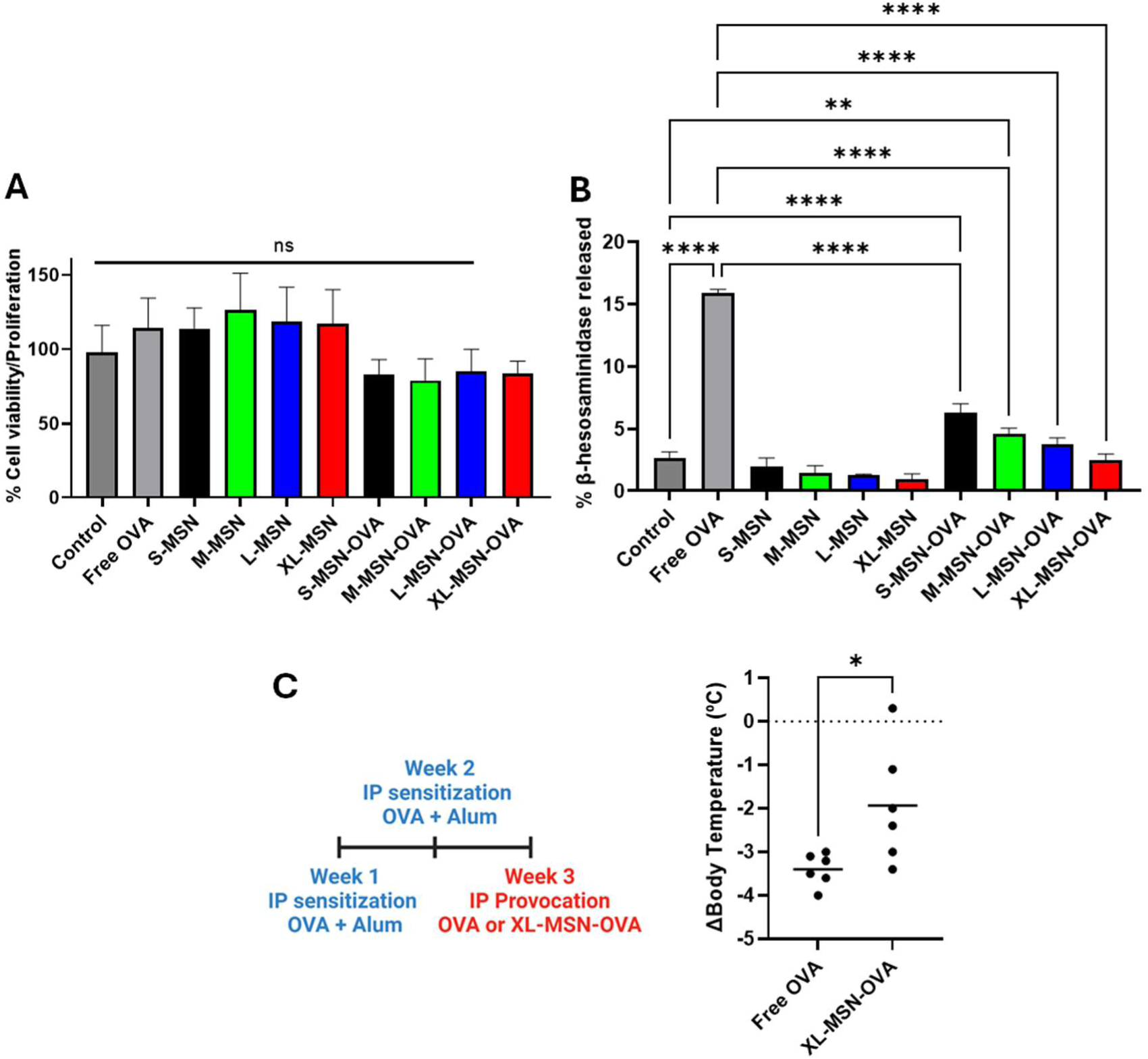
Safety evaluation of OVA-loaded MSNs. WST-1 cytotoxicity assay in RBL-2H3 cells (A). β-Hexosaminidase release assay in RBL-2H3 cells pre- sensitized with mouse anti-OVA IgE after incubation with free OVA, empty MSNs. and OVA-loaded MSNs, at a constant OVA concentration of 10 µg/mL (B). *In vivo* safety evaluation by measuring the body temperature drop in anaphylactic mice 20 minutes after the intraperitoneal injection of 500 µg of OVA, either free or loaded in XL-MSNs (C). n=6 (3 female, 3 male mice), statistical analysis performed by the Mann Whitney test, *p<0.05.

To further confirm this potential, an *in vivo* challenge was conducted by intraperitoneally injecting a large dose of OVA (500 µg) either free or loaded inside XL-MSNs to OVA-anaphylactic mice and evaluating the induction of systemic anaphylaxis reaction by measuring the body temperature drop (characteristic of anaphylaxis). As can be seen in **Figure 4C**, OVA loading inside XL-MSN led to a reduction in the body temperature drop after injection in anaphylactic mice compared to free protein. This result demonstrates that OVA loading in nanoparticles decreases the capacity of inducing an anaphylactic reaction *in vivo* compared to the same dose of free protein, confirming the improvement in the safety of this material as an AIT agent.

In AIT, the route of administration is of huge importance, as it determines which cells will be in contact with the therapeutic agent, influencing not only the safety of the therapy, but also its efficacy. Thus, to understand the potential effect of the route of administration on the performance of MSNs for AIT, their biodistribution after administration through different routes must first be assessed. To do this, the biodistribution of NIR-fluorescent MSNs was evaluated by an *in vivo* fluorescence imaging system after administration via the SC, SL, or IV route (**Figure 5**). It is worth noting that, as the fluorophore was introduced during nanoparticle synthesis, and is covalently bound to the silica network, the fluorescent signal detected corresponds to the presence of MSNs or degradation products of the nanoparticles, rather than any released dye from the particles. After SC administration, the fluorescence at the injection site decreases relatively fast, so that at 24 h after injection, only a small signal can be seen in the area (**Figure 5A**). On the other hand, a large fluorescence signal can be seen in the bladder at the 2 to 4 h time points, indicating the rapid elimination in urine of soluble degradation products. No relevant fluorescent signal could be detected in the extracted organs (liver, spleen, kidneys, tongue, lungs) 48 h after SC administration. After SL administration (**Figure 5B**), the fluorescent signal in the mouth was only detectable 1 h after administration, and up to 4 h after the fluorescent signal from the MSNs could be observed at different parts of the gastrointestinal tract, indicating a fast elimination of the MSNs following the swallowing of the sublingually administered nanoparticles. Consequently, no fluorescent signal could be seen in the extracted organs (including the tongue). Given these results, any observed effect from this route of administration would be difficult to distinguish from oral administration, as most (if not all) of the nanoparticle dose seems to be swallowed relatively fast and eliminated from the animals through the feces. After IV administration through tail vein injection (**Figure 5C**), intense fluorescence in the bladder could be observed at even shorter timepoints than after SC administration. In addition to this, further fluorescence could be detected in the upper half of the mouse, probably due to the presence of the MSNs in the pulmonary circulation. Interestingly, at longer time points, when fluorescence could no longer be detected in the bladder, a clear fluorescent signal could be seen in the abdominal region. This could be related to the accumulation of the administered MSNs in liver and/or spleen, as was later confirmed *ex vivo*. Given the difficulty in achieving reproducible tail vein injections when multiple administrations are needed, the retro-orbital IV route was also evaluated, observing equivalent biodistribution to tail vein injection (**Figure S2**). These biodistribution results are in good agreement with previous work reported by other groups. The biodegradation of MSNs into non-toxic soluble products has been widely reported [43,44], and silica degradation products had been previously detected in urine at time points as short as 30 min after IV injection [45]. On the other hand, the accumulation of nanoparticles in the liver and spleen after IV administration is well known in the nanomedicine field[46–49], and it is the main motivating factor for using nanoparticles for AIT [21], taking advantage of this passive accumulation of nanoparticles in these organs with large APC populations and, in the case of the liver, even with a tolerogenic environment[15,46,50]. Finally, regarding the equivalent biodistribution of IV administration through tail vein and retro-orbital injection, several studies have already reported that both injection methods could be used as interchangeably for different therapeutic agents, including antibodies and nanoparticles [51,52].

**Figure 5.**
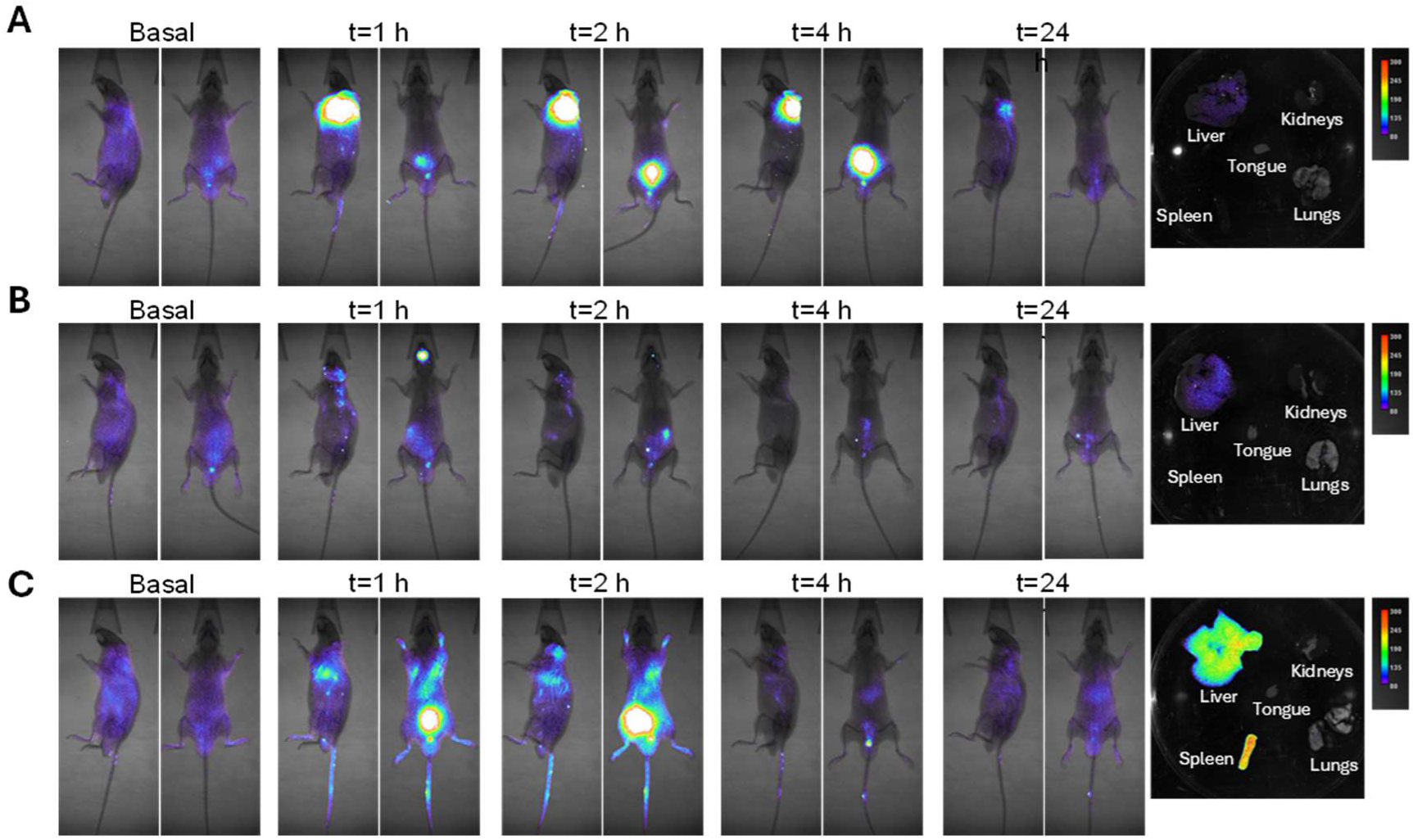
*In vivo* fluorescence images showing biodistribution of MSN at different time points after administration by subcutaneous (A), sublingual (B) or intravenous (tail vein injection, C) administration. *Ex vivo* fluorescence of different organs extracted 48 hours after MSN administration.

Finally, the therapeutic efficacy of OVA-loaded XL-MSNs for AIT was evaluated in a mouse model of systemic anaphylaxis to OVA (**Figure 6**). OVA-loaded XL- MSNs were administered weekly for 6 weeks via SC (SCIT), SL (SLIT), or IV (IVIT) routes to assess the impact of the administration route of therapeutic efficacy, considering the vastly different biodistribution profiles observed for each method **Figure 6A**). The results show that OVA-loaded XL-MSNs were only capable of reversing the anaphylactic state of the animals through IVIT, with all other experimental groups of sensitized mice (anaphylactic control, SCIT, and SLIT) showing no significant differences in body temperature drop upon intraperitoneal provocation with OVA (**Figure 6B**). This was further confirmed by the symptom scores of the animals after provocation **Figure 6C**), which only showed a reduction in the severity of the symptoms in the IVIT group compared to the anaphylactic control. Finally, all sensitized mice presented similar values of serum anti-OVA IgG1 and IgG2b (**Figure 6D-G**), but only the IVIT group showed an increase in the Th1-associated anti-OVA IgG2a, which correlates well with the therapeutic efficacy observed by the previous two parameters. It is worth noting that the SCIT group even exhibited higher levels of serum anti-OVA IgE.

**Figure 6.**
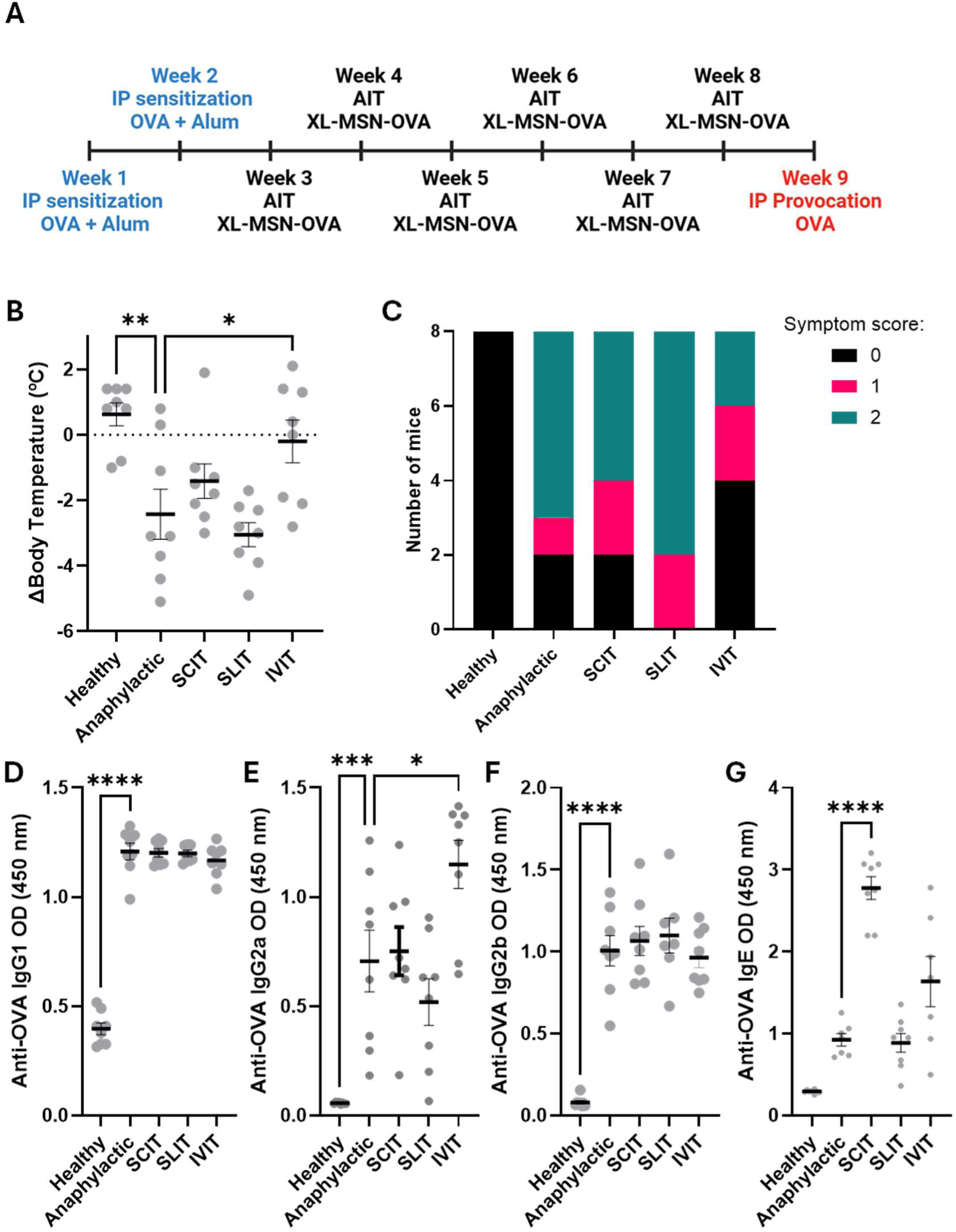
Therapeutic efficacy of OVA-loaded XL-MSNs for AIT in a mouse model of systemic anaphylaxis. Experimental timeline (A). Body temperature drop 20 minutes after provocation via intraperitoneal injection of OVA in healthy control, anaphylactic control, SCIT, SLIT, ant IVIT mice (B). Symptom score 20 minutes after provocation through intraperitoneal injection of OVA (C). Serum anti-OVA antibodies determined by ELISA after provocation: IgG1 (D), IgG2a (E), IgG2b (F) and IgE (G). Data are presented as individual values with means ± SEM (n = 8 per group). Statistical significance was determined by one-way ANOVA with appropriate post-hoc tests. *p<0.05; **p<0.01; ***p<0.005; ****p<0.001.

To further evaluate the immune changes induced by AIT with OVA-loaded XL- MSNs, we examined changes in different populations of the innate and adaptive immune systems in both the spleen and lymph nodes by flow cytometry. Anaphylactic mice did not show any differences in the frequency or phenotype of splenic dendritic cells compared to healthy (non-sensitized) controls or when AIT was administered via SCIT (**Figure 7A**). Nevertheless, IVIT or SLIT resulted in an increase in DCs (**Figure 7A**). Within the DC compartment, conventional DC1 (cDC1) and conventional DC2 (cDC2) subsets (**Figure 7B-C**) were more balanced in the immunotherapy-treated mice. This suggests a normalization of antigen presentation dynamics since cDC2s are responsible for presenting to CD4+ T cells, influencing class switching in B cells, and modulating allergic and inflammatory pathways. Similarly, plasmacytoid DCs (pDCs) (**Figure 7D**), often associated with regulatory processes, increased in the IVIT and SLIT groups, indicating a more tolerogenic DC status. Other innate immune populations, such as neutrophils and macrophages, were also examined (**Figure S1**), revealing more subtle differences, except for macrophages that resembled those of the DCs. The B cell compartment and plasma cells (PCs) demonstrated that immunotherapy treatment ameliorates the skewed humoral responses of the anaphylactic group in those mice receiving IVIT or SLIT (**Figure 7E-F**). Rather than sustaining high IgE production, MSN AIT appeared to support more controlled antibody production. We also observed an increase in Tregs (**Figure 7G**), which reduce allergic effector responses and promote peripheral tolerance. When splenocytes were cultured for 48 hours in the presence of OVA, IL-10 and IL-6 production increased in all sensitized groups compared to healthy controls, though not significantly, except for the IVIT group, which exhibited the highest cytokine levels (**Figure S1**). This observation adds to the evidence for a significant shift towards tolerance in the IVIT group.

**Figure 7.**
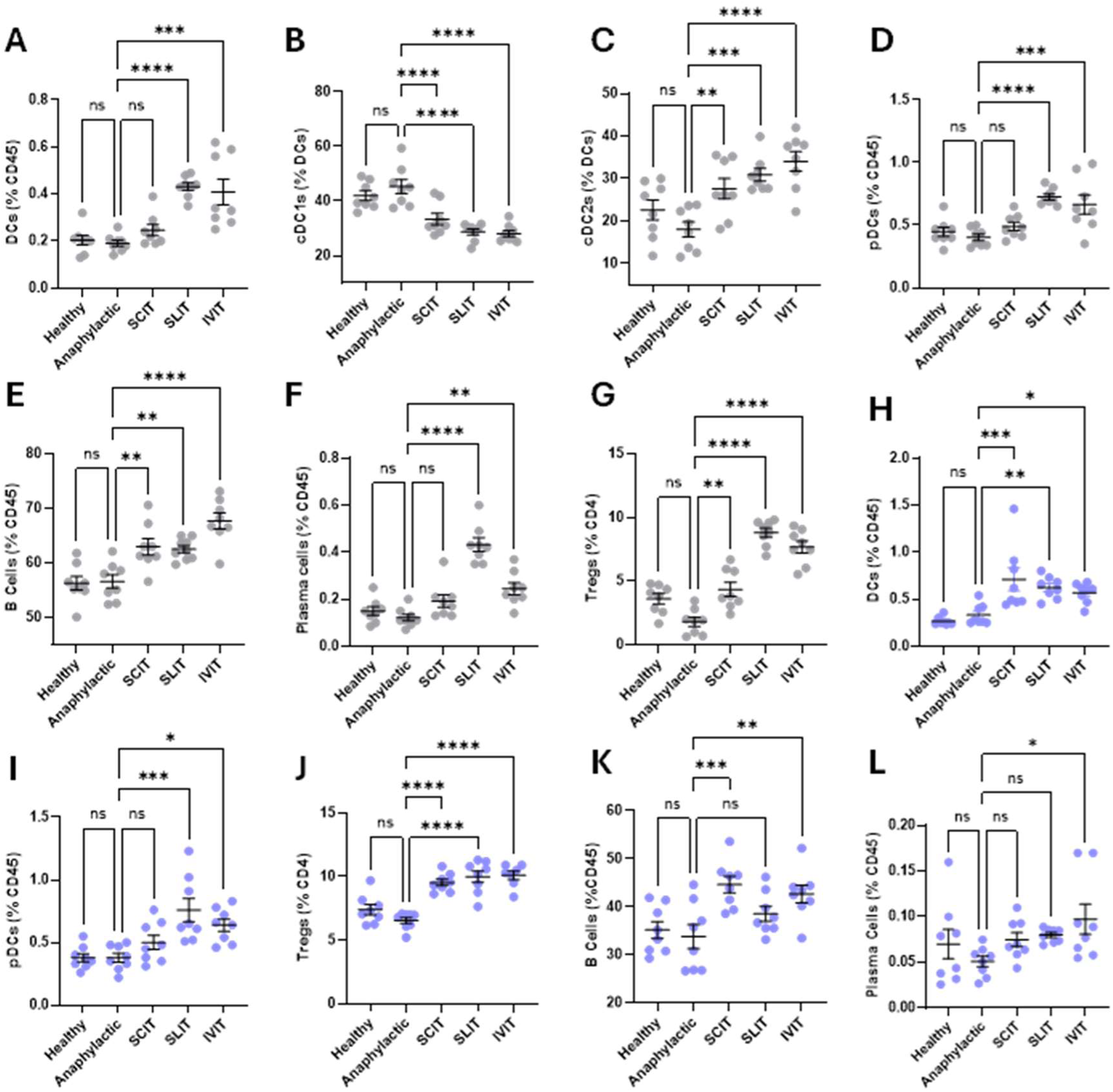
Flow cytometric analysis of immune cell populations isolated from spleens (grey, A-G) and lymph nodes (blue, H-L) under different experimental conditions: Healthy, Anaphylactic, subcutaneous immunotherapy (SCT), sublingual immunotherapy (SLIT), and intravenous immunotherapy (IVIT). Each panel shows the frequency of a specific immune cell subset, expressed as a percentage of the indicated parent population (e.g., %CD45 for total leukocytes, %DCs for dendritic cells, %CD4 for CD4^+ T cells). Total DC frequencies (A, H); cDC1 subsets (B); cDC2 subsets (C); pDC subsets (D, I); B cells (E, K); plasma cells (F, L); regulatory T cells (Tregs, G, J). Data are presented as individual values with means ± SEM (n = 8 per group). Statistical significance was determined by one-way ANOVA with appropriate post-hoc tests: ns = not significant; *p < 0.05; **p < 0.01; ***p < 0.005; ****p < 0.001.

Analysis of the draining lymph nodes showed immunological shifts toward tolerance in the animals receiving immunotherapy, which aligned with the patterns observed in the spleen. Total DC frequency rose noticeably compared to the anaphylaxis control, as did pDCs (**Figure 7H-I**). In parallel, Treg frequencies (**Figure 7J**) were significantly elevated in treated groups relative to the anaphylactic group, supporting the concept that immunotherapy enhances regulatory mechanisms that can dampen excessive allergic responses in peripheral lymphoid tissues. Additionally, both B cells and plasma cells increased in frequency compared to the anaphylactic group (**Figure 7K-L**). This suggests that the immunotherapy-induced modulation of the humoral compartment may not be simply related to the frequency in these populations, but rather involves a reshaping of the B cell response, including antibody isotype switching.

## 4. Conclusions

This work demonstrates the potential of MSNs as a promising platform for AIT. By employing mesoporous silica nanoparticles with a pore size optimized to load and release ovalbumin as a model allergen, their performance for AIT could be optimized, increasing their immunological capacity and suppressing the allergenicity compared to free protein. These immunogenic and allergenic potential was also confirmed *in vivo* before evaluating their biodistribution after subcutaneous, sublingual, and intravenous administration. Therapeutic evaluations in a mouse model of systemic anaphylaxis demonstrated that only intravenously administered OVA-loaded XL-MSNs successfully prevented anaphylaxis upon allergen provocation, associated with the production of ovalbumin-specific IgG2a (associated with a Th1 response) as well as to the increase in the Treg population, among other changes in the populations of the innate and adaptive immune system in both the spleen and lymph nodes. These findings underscore the potential of OVA-loaded XL-MSNs ovalbumin-loaded mesoporous silica nanoparticles as a safe and effective platform for AIT. Future work should explore modifications of these nanoparticles with targeting and/or adjuvant molecules to further improve their potential for this application.

## Acknowledgements

This work was supported by the Institute of Health “Carlos III” (ISCIII) (PI21/00346, RD21/0002/0008) cofunded by the European Union, and by the project PID2022-142781OA-I00 funded by MCIU/AEI/10.13039/501100011033/FEDER, UEU. APM acknowledges PhD fellowship PREP2022-000567 funded by MCIU/AEI/10.13039/501100011033 and by ESF+. PT, MCMA and JACL acknowledge Predoctoral Health Research Training Funding (FI21/00116, FI22/00106, FI22/00199) funded by ISCIII and by European Union NextGenerationEU/PRTR. CJA was supported by a Marie Sklodowska-Curie Actions (MSCA) Postdoctoral Fellowships from Horizon Europe (101105416). CM holds a “Nicolas Monardes” research contract by Andalusian Regional Ministry Health (RC0004-2021). JLP also acknowledges grant RYC2021-034536-I funded by MCIU/AEI/10.13039/501100011033 and by European Union NextGenerationEU/PRTR. TEM, fluorescence microscopy and *in vivo* imaging experiments were performed in the ICTS “NANBIOSIS,” more specifically in the U28 Unit at IBIMA Plataforma BIONAND.

## References

[1] D.D. Chaplin, Overview of the immune response, Journal of Allergy and Clinical Immunology 125 (2010) S3–S23. 10.1016/j.jaci.2009.12.980.

[2] M.I. Montañez, C. Mayorga, G. Bogas, E. Barrionuevo, R. Fernandez-Santamaria, A. Martin-Serrano, J.J. Laguna, M.J. Torres, T.D. Fernandez, I. Doña, Epidemiology, mechanisms, and diagnosis of drug-induced anaphylaxis, Front Immunol 8 (2017) 1–10. 10.3389/fimmu.2017.00614.

[3] B.T. Kelly, M.H. Grayson, Immunoglobulin E, what is it good for?, Annals of Allergy, Asthma and Immunology 116 (2016) 183–187. 10.1016/j.anai.2015.10.026.

[4] J.M. Brown, T.M. Wilson, D.D. Metcalfe, The mast cell and allergic diseases: role in pathogenesis and implications for therapy, Clinical & Experimental Allergy 38 (2008) 4–18. 10.1111/J.1365-2222.2007.02886.X.

[5] J.L. Paris, P. de la Torre, A.I. Flores, New Therapeutic Approaches for Allergy: A Review of Cell Therapy and Bio- or Nano-Material-Based Strategies, Pharmaceutics 13 (2021) 2149. 10.3390/pharmaceutics13122149.

[6] M.J. Rodriguez, A. Mascaraque, J. Ramos-Soriano, M.J. Torres, J.R. Perkins, F. Gomez, M. Garrido-Arandia, N. Cubells-Baeza, D. Andreu, A. Diaz-Perales, J. Rojo, C. Mayorga, Pru p 3-Epitope-based sublingual immunotherapy in a murine model for the treatment of peach allergy, Mol Nutr Food Res 61 (2017) 1700110. 10.1002/mnfr.201700110.

[7] L. Johnson, A. Duschl, M. Himly, Nanotechnology-Based Vaccines for Allergen-Specific Immunotherapy: Potentials and Challenges of Conventional and Novel Adjuvants under Research, Vaccines (Basel) 8 (2020) 237. 10.3390/vaccines8020237.

[8] J. De Souza Rebouças, I. Esparza, M. Ferrer, M.L. Sanz, J.M. Irache, C. Gamazo, Nanoparticulate adjuvants and delivery systems for allergen immunotherapy, J Biomed Biotechnol 2012 (2012). 10.1155/2012/474605.

[9] C. Gamazo, G. Gastaminza, M. Ferrer, M.L. Sanz, J.M. Irache, Nanoparticle based-immunotherapy against allergy, Immunotherapy 6 (2014) 885–897. 10.2217/imt.14.63.

[10] J.L. Paris, A. Baeza, Nano- and Microscale Drug Delivery Approaches for Therapeutic Immunomodulation, ChemNanoMat 7 (2021) 773–788. 10.1002/cnma.202100062.

[11] F. Palomares, J. Ramos-Soriano, F. Gomez, A. Mascaraque, G. Bogas, J.R. Perkins, M. Gonzalez, M.J. Torres, A. Diaz-Perales, J. Rojo, C. Mayorga, Pru p 3-Glycodendropeptides Based on Mannoses Promote Changes in the Immunological Properties of Dendritic and T-Cells from LTP-Allergic Patients, Mol Nutr Food Res 63 (2019) 1–11. 10.1002/mnfr.201900553.

[12] K.R. Hughes, M.N. Saunders, J.J. Landers, K.W. Janczak, H. Turkistani, L.M. Rad, S.D. Miller, J.R. Podojil, L.D. Shea, J.J. O’Konek, Masked Delivery of Allergen in Nanoparticles Safely Attenuates Anaphylactic Response in Murine Models of Peanut Allergy, Frontiers in Allergy 3 (2022) 1–11. 10.3389/falgy.2022.829605.

[13] J.L. Paris, L.K. Vora, M.J. Torres, C. Mayorga, R.F. Donnelly, Microneedle array patches for allergen-specific immunotherapy, Drug Discov Today 28 (2023) 103556. 10.1016/j.drudis.2023.103556.

[14] P.Y. Li, F. Bearoff, P. Zhu, Z. Fan, Y. Zhu, M. Fan, L. Cort, T. Kambayashi, E.P. Blankenhorn, H. Cheng, PEGylation enables subcutaneously administered nanoparticles to induce antigen-specific immune tolerance, Journal of Controlled Release (2021) 135907. 10.1016/j.jconrel.2021.01.013.

[15] Q. Liu, X. Wang, X. Liu, S. Kumar, G. Gochman, Y. Ji, Y.-P. Liao, C.H. Chang, W. Situ, J. Lu, J. Jiang, K.-C. Mei, H. Meng, T. Xia, A.E. Nel, Use of Polymeric Nanoparticle Platform Targeting the Liver To Induce Treg-Mediated Antigen-Specific Immune Tolerance in a Pulmonary Allergen Sensitization Model, ACS Nano 13 (2019) 4778–4794. 10.1021/acsnano.9b01444.

[16] Q. Liu, X. Wang, X. Liu, Y.-P. Liao, C.H. Chang, K.-C. Mei, J. Jiang, S. Tseng, G. Gochman, M. Huang, Z. Thatcher, J. Li, S.D. Allen, L. Lucido, T. Xia, A.E. Nel, Antigen- and Epitope-Delivering Nanoparticles Targeting Liver Induce Comparable Immunotolerance in Allergic Airway Disease and Anaphylaxis as Nanoparticle-Delivering Pharmaceuticals, ACS Nano 15 (2021) 1608–1626. 10.1021/acsnano.0c09206.

[17] M.N. Saunders, L.M. Rad, L.A. Williams, J.J. Landers, R.R. Urie, S.E. Hocevar, M. Quiros, M. Chiang, A.R. Angadi, K.W. Janczak, E.J. Bealer, K. Crumley, O.E. Benson, K. V. Griffin, B.C. Ross, C.A. Parkos, A. Nusrat, S.D. Miller, J.R. Podojil, J.J. O’Konek, L.D. Shea, Allergen-Encapsulating Nanoparticles Reprogram Pathogenic Allergen-Specific Th2 Cells to Suppress Food Allergy, Adv Healthc Mater (2024). 10.1002/adhm.202400237.

[18] M.J. Rodriguez, J. Ramos-Soriano, J.R. Perkins, A. Mascaraque, M.J. Torres, F. Gomez, A. Diaz-Perales, J. Rojo, C. Mayorga, Glycosylated nanostructures in sublingual immunotherapy induce long-lasting tolerance in LTP allergy mouse model, Sci Rep 9 (2019) 4043. 10.1038/s41598-019-40114-7.

[19] A. Le Moignic, V. Malard, T. Benvegnu, L. Lemiègre, M. Berchel, P.A. Jaffrès, C. Baillou, M. Delost, R. Macedo, J. Rochefort, G. Lescaille, C. Pichon, F.M. Lemoine, P. Midoux, V. Mateo, Preclinical evaluation of mRNA trimannosylated lipopolyplexes as therapeutic cancer vaccines targeting dendritic cells, Journal of Controlled Release 278 (2018) 110–121. 10.1016/j.jconrel.2018.03.035.

[20] X. Le Guével, M. Perez Perrino, T.D. Fernández, F. Palomares, M.J. Torres, M. Blanca, J. Rojo, C. Mayorga, Multivalent Glycosylation of Fluorescent Gold Nanoclusters Promotes Increased Human Dendritic Cell Targeting via Multiple Endocytic Pathways, ACS Appl Mater Interfaces 7 (2015) 20945– 20956. 10.1021/acsami.5b06541.

[21] Q. Liu, X. Wang, Y.-P. Liao, C.H. Chang, J. Li, T. Xia, A.E. Nel, Use of a liver-targeting nanoparticle platform to intervene in peanut-induced anaphylaxis through delivery of an Ara h2 T-cell epitope, Nano Today 42 (2022) 101370. 10.1016/j.nantod.2021.101370.

[22] S.C. Balmert, C. Donahue, J.R. Vu, G. Erdos, L.D. Falo, S.R. Little, In vivo induction of regulatory T cells promotes allergen tolerance and suppresses allergic contact dermatitis, Journal of Controlled Release 261 (2017) 223–233. 10.1016/j.jconrel.2017.07.006.

[23] S. Shahgordi, M. Sankian, Y. Yazdani, K. Mashayekhi, S. Hasan Ayati, M. Sadeghi, M. Saeidi, M. Hashemi, Immune responses modulation by curcumin and allergen encapsulated into PLGA nanoparticles in mice model of rhinitis allergic through sublingual immunotherapy, Int Immunopharmacol 84 (2020) 106525. 10.1016/j.intimp.2020.106525.

[24] W.R. Reisacher, D. Liotta, The use of poly(D,L-lactic-co-glycolic) acid microspheres in the treatment of allergic disease, Curr Opin Otolaryngol Head Neck Surg 19 (2011) 188–192. 10.1097/MOO.0b013e328345013a.

[25] S. Gómez, C. Gamazo, B. San Roman, M. Ferrer, M.L. Sanz, S. Espuelas, J.M. Irache, Allergen immunotherapy with nanoparticles containing lipopolysaccharide from Brucella ovis, European Journal of Pharmaceutics and Biopharmaceutics 70 (2008) 711–717. 10.1016/j.ejpb.2008.05.016.

[26] S. Gómez, C. Gamazo, B.S. Roman, M. Ferrer, M.L. Sanz, J.M. Irache, Gantrez® AN nanoparticles as an adjuvant for oral immunotherapy with allergens, Vaccine 25 (2007) 5263–5271. 10.1016/j.vaccine.2007.05.020.

[27] H. Pohlit, I. Bellinghausen, H. Frey, J. Saloga, Recent advances in the use of nanoparticles for allergen-specific immunotherapy, Allergy: European Journal of Allergy and Clinical Immunology 72 (2017) 1461–1474. 10.1111/all.13199.

[28] S. Wu, Y. Hung, C. Mou, Mesoporous silica nanoparticles as nanocarriers, Chemical Communications 47 (2011) 9972–85. 10.1039/c1cc11760b.

[29] M. Manzano, M. Vallet-Regí, Mesoporous silica nanoparticles in nanomedicine applications, J Mater Sci Mater Med 29 (2018) 65. 10.1007/s10856-018-6069-x.

[30] X. Peng, Y. Liang, Y. Yin, H. Liao, L. Li, Development of a hollow mesoporous silica nanoparticles vaccine to protect against house dust mite induced allergic inflammation, Int J Pharm 549 (2018) 115–123. 10.1016/j.ijpharm.2018.07.047.

[31] J.L. Paris, M.V. Cabañas, M. Manzano, M. Vallet-Regí, Polymer-Grafted Mesoporous Silica Nanoparticles as Ultrasound-Responsive Drug Carriers, ACS Nano 9 (2015) 11023–11033. 10.1021/acsnano.5b04378.

[32] J.L. Paris, M. Colilla, I. Izquierdo-Barba, M. Manzano, M. Vallet-Regí, Tuning mesoporous silica dissolution in physiological environments: a review, J Mater Sci 52 (2017) 8761–8771. 10.1007/s10853-017-0787-1.

[33] J.L. Paris, C. Mannaris, M.V. Cabañas, R. Carlisle, M. Manzano, M. Vallet-Regí, C.C. Coussios, Ultrasound-mediated cavitation-enhanced extravasation of mesoporous silica nanoparticles for controlled-release drug delivery, Chemical Engineering Journal 340 (2018) 2–8. 10.1016/j.cej.2017.12.051.

[34] A.M. Pérez-Moreno, C.J. Aranda, M.J. Torres, C. Mayorga, J.L. Paris, Immunomodulatory potential of rapamycin-loaded mesoporous silica nanoparticles: pore size-dependent drug loading, release, and in vitro cellular responses, Drug Deliv Transl Res 14 (2024) 3467–3476. 10.1007/s13346-024-01575-0.

[35] J.L. Paris, L.K. Vora, A.M. Pérez-Moreno, M. del C. Martín-Astorga, Y.A. Naser, Q.K. Anjani, J.A. Cañas, M.J. Torres, C. Mayorga, R.F. Donnelly, Dissolving microneedle array patches containing mesoporous silica nanoparticles of different pore sizes as a tunable sustained release platform, Int J Pharm 669 (2025) 125064. 10.1016/j.ijpharm.2024.125064.

[36] X. Wang, X. Li, A. Ito, Y. Sogo, T. Ohno, Pore size-dependent immunogenic activity of mesoporous silica-based adjuvants in cancer immunotherapy, J Biomed Mater Res A 102 (2014) 967–974. 10.1002/jbm.a.34783.

[37] X. Hong, X. Zhong, G. Du, Y. Hou, Y. Zhang, Z. Zhang, T. Gong, L. Zhang, X. Sun, The pore size of mesoporous silica nanoparticles regulates their antigen delivery efficiency., Sci Adv 6 (2020) eaaz4462. 10.1126/sciadv.aaz4462.

[38] N. Kupferschmidt, K.R. Qazi, C. Kemi, H. Vallhov, A.E. Garcia-Bennett, S. Gabrielsson, A. Scheynius, Mesoporous silica particles potentiate antigen-specific T-cell responses, Nanomedicine 9 (2014) 1835–1846. 10.2217/NNM.13.170.

[39] P.N. Durfee, Y.-S. Lin, D.R. Dunphy, A.J. Muñiz, K.S. Butler, K.R. Humphrey, A.J. Lokke, J.O. Agola, S.S. Chou, I.-M. Chen, W. Wharton, J.L. Townson, C.L. Willman, C.J. Brinker, Mesoporous Silica Nanoparticle-Supported Lipid Bilayers (Protocells) for Active Targeting and Delivery to Individual Leukemia Cells, ACS Nano 10 (2016) 8325–8345. 10.1021/acsnano.6b02819.

[40] J.L. Paris, C. Monío, A.M. Pérez-Moreno, R. Jurado-Escobar, G. Bogas, T.D. Fernández, M.I. Montañez, C. Mayorga, M.J. Torres, Influence of Pore Size in Protein G’-Grafted Mesoporous Silica Nanoparticles as a Serum Pretreatment System for In Vitro Allergy Diagnosis, Adv Healthc Mater 2203321 (2023) 2203321. 10.1002/adhm.202203321.

[41] K.S. Butler, P.N. Durfee, C. Theron, C.E. Ashley, E.C. Carnes, C.J. Brinker, Protocells: Modular Mesoporous Silica Nanoparticle-Supported Lipid Bilayers for Drug Delivery, Small 12 (2016) 2173–2185. 10.1002/smll.201502119.

[42] B.G. Cha, J.H. Jeong, J. Kim, Extra-Large Pore Mesoporous Silica Nanoparticles Enabling Co-Delivery of High Amounts of Protein Antigen and Toll-like Receptor 9 Agonist for Enhanced Cancer Vaccine Efficacy, ACS Cent Sci 4 (2018) 484–492. 10.1021/acscentsci.8b00035.

[43] J.L. Paris, M. Colilla, I. Izquierdo-Barba, M. Manzano, M. Vallet-Regí, Tuning mesoporous silica dissolution in physiological environments: a review, J Mater Sci 52 (2017) 8761–8771. 10.1007/s10853-017-0787-1.

[44] K. Braun, A. Pochert, M. Beck, R. Fiedler, J. Gruber, M. Lindén, Dissolution kinetics of mesoporous silica nanoparticles in different simulated body fluids, J Solgel Sci Technol 79 (2016) 319–327. 10.1007/s10971-016-4053-9.

[45] Q. He, Z. Zhang, F. Gao, Y. Li, J. Shi, In vivo Biodistribution and Urinary Excretion of Mesoporous Silica Nanoparticles: Effects of Particle Size and PEGylation, Small 7 (2011) 271–280. 10.1002/smll.201001459.

[46] A. Carambia, B. Freund, D. Schwinge, O.T. Bruns, S.C. Salmen, H. Ittrich, R. Reimer, M. Heine, S. Huber, C. Waurisch, A. Eychmüller, D.C. Wraith, T. Korn, P. Nielsen, H. Weller, C. Schramm, S. Lüth, A.W. Lohse, J. Heeren, J. Herkel, Nanoparticle-based autoantigen delivery to Treg-inducing liver sinusoidal endothelial cells enables control of autoimmunity in mice, J Hepatol 62 (2015) 1349–1356. 10.1016/j.jhep.2015.01.006.

[47] L. García, M. Buñuales, N. Düzgüneş, C. Tros de Ilarduya, Serum-resistant lipopolyplexes for gene delivery to liver tumour cells, European Journal of Pharmaceutics and Biopharmaceutics 67 (2007) 58–66. 10.1016/j.ejpb.2007.01.005.

[48] S. Tammam, S. Mathur, N. Afifi, Preparation and Biopharmaceutical Evaluation of Tacrolimus Loaded Biodegradable Nanoparticles for Liver Targeting, J Biomed Nanotechnol 8 (2012) 439–449. 10.1166/jbn.2012.1403.

[49] S.K. Misra, G. Ghoshal, M.R. Gartia, Z. Wu, A.K. De, M. Ye, C.R. Bromfield, E.M. Williams, K. Singh, K. V Tangella, L. Rund, K. Schulten, L.B. Schook, P.S. Ray, E.C. Burdette, D. Pan, Trimodal Therapy: Combining Hyperthermia with Repurposed Bexarotene and Ultrasound for Treating Liver Cancer., ACS Nano 9 (2015) 10695–718. 10.1021/acsnano.5b05974.

[50] A. Carambia, C. Gottwick, D. Schwinge, S. Stein, R. Digigow, M. Şeleci, D. Mungalpara, M. Heine, F.A. Schuran, C. Corban, A.W. Lohse, C. Schramm, J. Heeren, J. Herkel, Nanoparticle-mediated targeting of autoantigen peptide to cross-presenting liver sinusoidal endothelial cells protects from CD8 T-cell-driven autoimmune cholangitis, Immunology (2021) imm.13298. 10.1111/imm.13298.

[51] A. Schoch, I.S. Thorey, J. Engert, G. Winter, T. Emrich, Comparison of the lateral tail vein and the retro-orbital venous sinus routes of antibody administration in pharmacokinetic studies, Lab Anim (NY) 43 (2014) 95–99. 10.1038/laban.481.

[52] M. Rouchota, A. Adamiano, M. Iafisco, E. Fragogeorgi, I. Pilatis, G. Doumont, S. Boutry, D. Catalucci, A. Zacharioudaki, G.C. Kagadis, Optimization of In Vivo Studies by Combining Planar Dynamic and Tomographic Imaging: Workflow Evaluation on a Superparamagnetic Nanoparticles System, Mol Imaging 2021 (2021). 10.1155/2021/6677847.

