## Supporting Information for "Ovalbumin-loaded mesoporous silica nanoparticles for allergen specific immunotherapy"

for

**Table S1**. Antibodies for flow cytometry employed in this work.

| **Molecule** | **Fluorochrome** | **Catalog #** | **Manufacturer** | **Clone** | **Dilution** |
|---|---|---|---|---|---|
| CD138 | BV421 | 142523 | Biolegend | 281-2 | 100 |
| CD40 | PB | 124625 | Biolegend | 3/23 | 50 |
| CD25 | BV480 | 566120 | BD Biosciences | PC61 | 50 |
| CD103 | BV510 | 563087 | BD Biosciences | M290 | 50 |
| CD11b | BV605 | 101257 | Biolegend | M1/70 | 200 |
| CD83 | BV650 | 121515 | Biolegend | Michel-19 | 50 |
| MHC-II | BV711 | 107643 | Biolegend | M5/114.15.2 | 400 |
| CD3 | BV750 | 100249 | Biolegend | 17A2 | 100 |
| B220 | BV785 | 103245 | Biolegend | RA3-6B2 | 200 |

| | | | | | |
|---|---|---|---|---|---|
| Foxp3 | AF488 | 126405 | Biolegend | MF-14 | 50 |
| CD4 | Spark Blue 550 | 100473 | Biolegend | GK1.5 | 800 |
| Ly-6C | PE-Dazzle 594 | 128043 | Biolegend | HK1.4 | 400 |
| CD80 | PE-Fire 640 | 104759 | Biolegend | 16-10A1 | 100 |
| F4/80 | RB705 | 570289 | BD Biosciences | T45-2342 | 400 |
| CD8α | RB744 | 570486 | BD Biosciences | 53-6.7 | 400 |
| CD11c | PE- Fire 810 | 161105 | Biolegend | N418 | 200 |
| CD86 | APC-Cy7 | 105029 | Biolegend | GL1 | 50 |
| NK1.1 | APC-Fire 810 | 156520 | Biolegend | S17016D | 400 |
| p-DCA | Spark UV 387 | 127043 | Biolegend | 927 | 150 |
| CD45 | BUV661 | 612975 | BD Biosciences | 30-F11 | 400 |
| Ly-6G | BUV737 | 568346 | BD Biosciences | 1A8 | 200 |
| Ki-67 | BUV805 | 368-5698-82 | Invitrogen | SolA15 | 400 |

**Table S2**. Characterization of the prepared MSNs.

| | S-MSN | M-MSN | L-MSN | XL-MSN |
|---|---|---|---|---|
| Hydrodynamic diameter (Z Average, nm) | 157.06 ± 1.58 | 165.27 ± 2.00 | 168.85 ± 5.10 | 161.73 ± 3.32 |
| PDI | 0.21 ± 0.02 | 0.34 ± 0.01 | 0.30 ± 0.06 | 0.21 ± 0.01 |
| Z Potential (mV) | -20.4 ± 0.4 | -23.1 ± 0.8 | -13.7 ± 0.5 | -18.7 ± 0.5 |
| BET* Surface Area ($m^2/g$) | 288.6 | 921.9 | 587.1 | 971.4 |
| Pore volume ($cm^3/g$) | 0.35 | 1.35 | 0.96 | 2.04 |
| Adsorption average pore diameter (nm) | 4.85 | 5.85 | 6.51 | 8.39 |

**Table S3**. *In vivo* OVA-loaded XL-MSN biocompatibility after 3 weekly subcutaneous injections (serum biomarkers).

| | Control | MSN-OVA SC |
|---|---|---|
| Albumin (g/dL) | 4.66 ± 1.12 | 4.77 ± 0.28 |
| LDH (U/L) | 92.83 ± 33.33 | 104.21 ± 3.82 |
| Total Protein (g/dL) | 6.29 ± 0.78 | 6.45 ± 0.27 |
| SGPT (U/L) | 18.40 ± 10.28 | 13.03 ± 3.99 |
| AST/SGOT (U/L) | 106.25 ± 5.01 | 118.33 ± 14.27 |

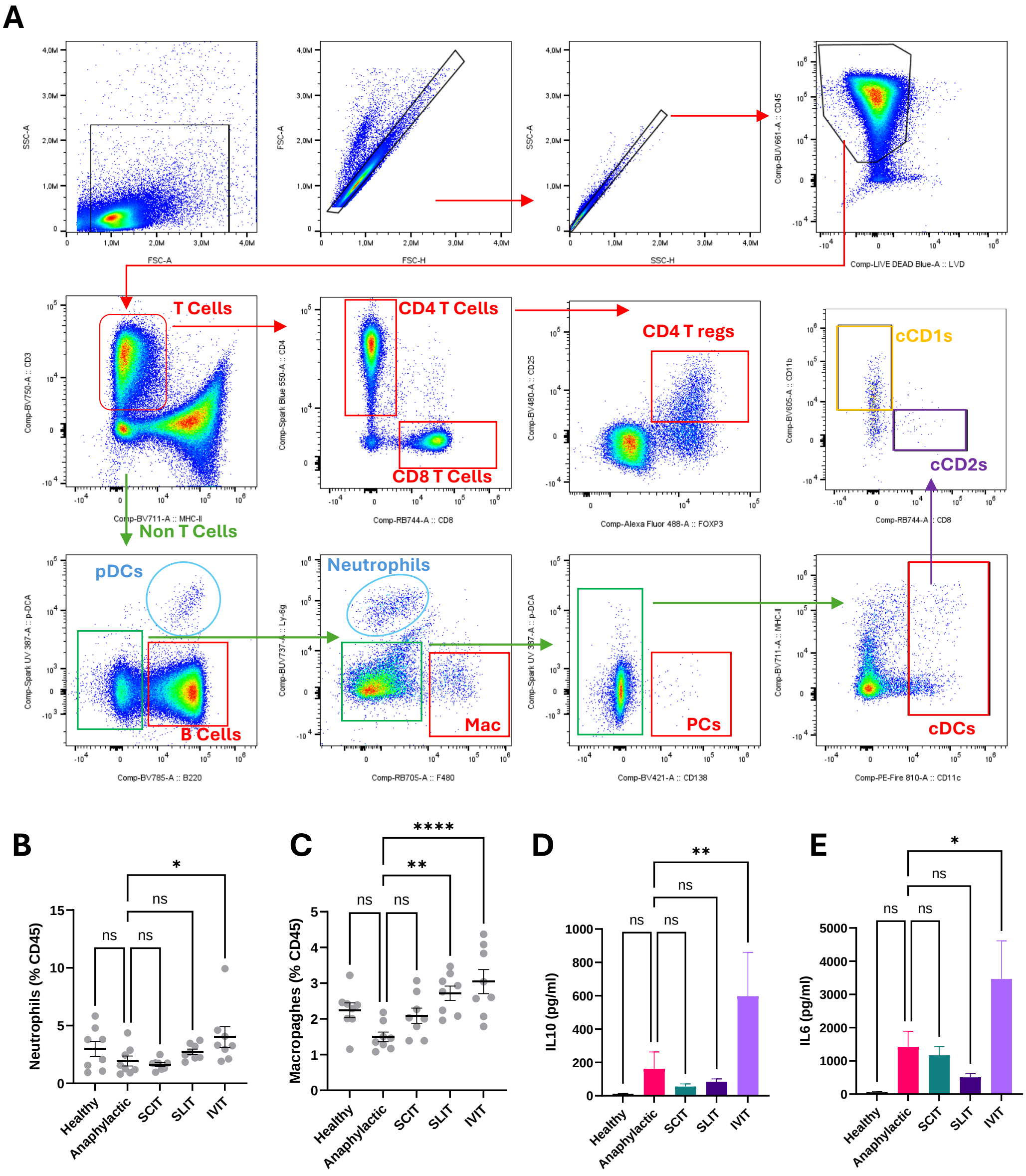


**Figure S1.** Gating strategy for flow cytometry of spleen and lymph node cells (A). Evaluation of neutrophil (B) and macrophage (C) populations as function of CD45+ cells in spleen. Production of IL-10 (D) and IL-6 (E) by splenocytes after cell culture for 48 h. Data are Means ±SEM, n=8. Statistical analysis by One way ANOVA. *p<0.05; **p<0.01; ****p<0.001.

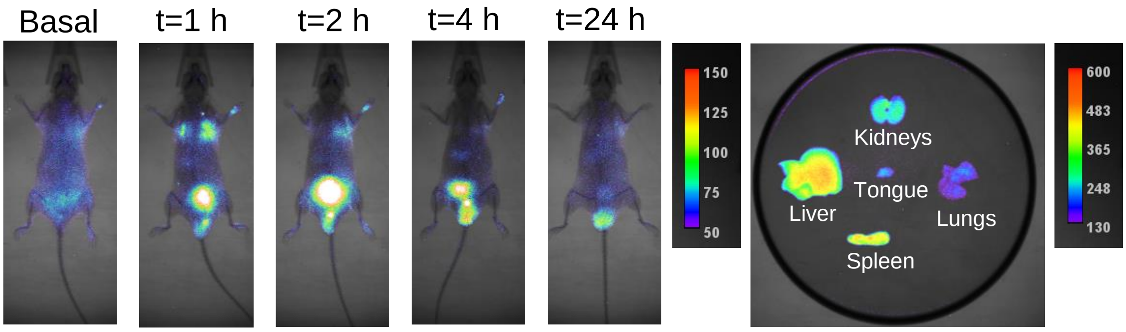


**Figure S2.** *In vivo* fluorescence images showing biodistribution of MSN at different timepoints after intravenous administration by retro-orbital injection. *Ex vivo* fluorescence of different organs extracted 24 h after MSN administration.
